# Morphometric analysis of *Passiflora* leaves: the relationship between landmarks of the vasculature and elliptical Fourier descriptors of the blade

**DOI:** 10.1101/067512

**Authors:** Daniel H. Chitwood, Wagner C. Otoni

## Abstract

**BACKGROUND:** Leaf shape among *Passiflora* species is spectacularly diverse. Underlying this diversity in leaf shape are profound changes in the patterning of the primary vasculature and laminar outgrowth. Each of these aspects of leaf morphology—vasculature and blade—provides different insights into leaf patterning.

**RESULTS:** Here, we morphometrically analyze >3,300 leaves from 40 different *Passiflora* species collected sequentially across the vine. Each leaf is measured in two different ways: using 1) 15 homologous Procrustes-adjusted landmarks of the vasculature, sinuses, and lobes and 2) Elliptical Fourier Descriptors (EFDs), which quantify the outline of the leaf. The ability of landmarks, EFDs, and both datasets together are compared to determine their relative ability to predict species and node position within the vine. Pairwise correlation of x and y landmark coordinates and EFD harmonic coefficients reveals close associations between traits and insights into the relationship between vasculature and blade patterning.

**CONCLUSIONS:** Landmarks, more reflective of the vasculature, and EFDs, more reflective of the blade contour, describe both similar and distinct features of leaf morphology. Landmarks and EFDs vary in ability to predict species identity and node position in the vine and exhibit a correlational structure (both within landmark or EFD traits and between the two data types) revealing constraints between vascular and blade patterning underlying natural variation in leaf morphology among *Passiflora* species.

## Background

The leaves of *Passiflora* species are remarkably diverse [1–3]. The underlying source of such diversity is ultimately speculative, but diversifying selective pressure from egg-laying *Heliconius* butterflies that use leaf shape as a visual cue has been proposed [4–6]. The leaves not only vary between species, but between successive nodes of a single vine, sometimes dramatically, reflecting both the heteroblastic development of the shoot apical meristem from which they are derived and the ontogeny of individual leaves as they allometrically expand [7–10]. Previous morphometric work using the multiscale Minkowski fractal dimension focused on vein patterning and the contour of the blade to predictively identify *Passiflora* species. Of the 10 species analyzed, some possessed similar leaf morphologies that could be correctly classified using only a small number of leaves per species as a training set [11].

To some degree, the patterning of the vein and blade follow each other, but to what degree they vary independently, or one is the consequence of the other, remains to be determined [12–15]. At a morphometric level, vascular patterning and the contour of the blade are studied separately, as one is a topology and the other a contour. Vasculature and blade can be separated and then analyzed with the same method, and was done using a Fractal-based approach in *Passiflora* previously [11].

Alternatively, traditional morphometric approaches can be applied to vascular patterning and the outline of the blade [16]. Procrustes-adjusted landmarks are coordinate points that correspond between all measured samples, ideally through homology [17]. Homologous landmarks are ideally suited for measuring vein thickness, vascular branch points, and the relative positioning and depth of sinuses and lobes if these features can be found in every sample, as in many palmately-lobed species, such as *Cucurbita, Acer,* and *Vitis.* [10, 18–25].

The landmark concept can be applied to contours as well, placing numerous points along a curve and subsequently using a Procrustes superimposition to create a near-continuous analysis of outlines [26]. The pseudo-landmark approach to quantifying contours has been used extensively to study leaf outlines, especially in species like *Antirrhinum* and *Arabidopsis,* where homologous points are lacking [27–29]. Another approach, Elliptical Fourier Descriptors (EFDs), treats an outline as a wave connecting back onto itself and subsequently performs a Fourier transform, decomposing the shape into a harmonic series [30–33]. EFDs have been applied to species from both *Solanum* and *Vitis* [23, 34–38] as well as the study of leaf asymmetry [25, 39, 40], leveraging the ability to separate symmetric from asymmetric sources of variance.

Comparing landmark- and contour-based methods not only provides an integrated perspective on leaf morphology, but can also potentially reveal the extent that patterning of the vasculature and blade are correlated in a quantitative fashion. Understanding the complementary features different morphometric methods detect is relevant to a wide variety of fields that use different approaches to extract information content from leaf shapes, including paleobiology and paleoclimate studies [41], ecology [42], evolution [10, 24, 27, 34, 43], genetics [21, 23, 29, 35, 36, 38], developmental biology [10, 18, 20, 24, 25, 34, 36, 39, 40], and plasticity [19, 20, 24, 37]. *Heliconius* butterflies, too, can even distinguish the shapes of leaves from different *Passiflora* species, presumably using a learning method yet to be determined [6].

Here, we measure landmarks of the vasculature, sinuses, and lobes and EFDs of the blade for >3,300 leaves from 40 *Passiflora* species sampled from successive nodes across vines. Linear Discriminant Analyses (LDAs) are used to determine the capacity of landmarks, EFDs, or both datasets to predict species identity versus node position in the vine. A correlational analysis of landmark and EFD data determines which specific features of leaves change together versus vary independently from each other. Our data reveals the constraints between vascular and blade patterning underlying natural variation in leaf morphology among *Passiflora* species and provides a critical comparison of complementary morphometric approaches used on the same leaves.

## Data Description

The purpose of this manuscript is to compare and contrast landmark and Elliptical Fourier Descriptor (EFD) methods in the genetic and developmental analysis of leaf shape among *Passiflora* species across the sequential nodes of their vines. The dataset released with this manuscript [44] consists of 555 scans of leaves from 40 different species of *Passiflora* in which the order of leaves arising from the vine is recorded, starting with “1” for the youngest leaf scanned from the growing tip of each vine. We importantly note: the numbering of nodes in the raw scans described above, starting at the tip of the shoot, is opposite from the numbering of nodes presented in the manuscript, in which numbering (starting with “1”) begins with the oldest leaf at the base of the shoot. The reason for this opposite numbering in the manuscript is that by beginning the counting of nodes with “1” at the shoot base the numbering aligns with the heteroblastic series (which begins with the first emerged leaf at the shoot base). >3,300 leaves are represented in this dataset. The number of vines sampled per a species and the number of nodes sampled for each vine are indicated in the raw data provided with this manuscript [45] and are visually depicted as well (**Fig. S1**). Both landmark data, measuring the vasculature, lobes, and sinuses, and Elliptical Fourier Descriptor (EFD) data, which quantify the leaf outline, can be derived from the provided datasets. Code used in the statistical analysis of data is also provided [45]. It is hoped that the release of this data will assist others in developing novel morphometric approaches to better understand the genetic, developmental, and environmental basis of leaf shape.

## Analyses

### Vascular landmarks and Elliptical Fourier Descriptors (EFDs) of the blade

For the >3,300 leaves measured across the leaf series for 40 different *Passiflora* species, a comparison of homologous landmarks and Elliptical Fourier Descriptors (EFDs) was made (see **Fig. S1** and raw data [45] for the replication associated with each species and the number of nodes for each vine) These two methods globally capture complementary aspects of leaf shape, sensitive to vascular patterning and the shape of the blade, respectively.

15 landmarks were measured for each leaf (Fig. 1A). For the proximal veins (near the leaf base) landmarks on each side of the junction of the proximal vein with the petiolar junction (where the major veins meet) were placed (landmarks 1-2 and 5-6), capturing the width of the proximal veins. Landmarks placed at the tip of the proximal vein (landmarks 7 and 15) capture the length and angle of the proximal lobe. On the distal vein (nearer the leaf tip), landmarks were placed only on the distal side of the junction with the midvein (landmarks 3 and 4) as the other side of the base of the distal vein variably intersects the midvein, petiolar junction, and proximal vein (see three examples in Fig. 1A, left to right). The landmarks at the tip of the distal veins (landmarks 13 and 9) measure the length and angle of the distal lobe. Additionally, landmarks describe the placement of the leaf tip (landmark 11), distal sinuses (landmarks 10 and 12), and the proximal sinuses (8 and 14).

**Figure 1:**
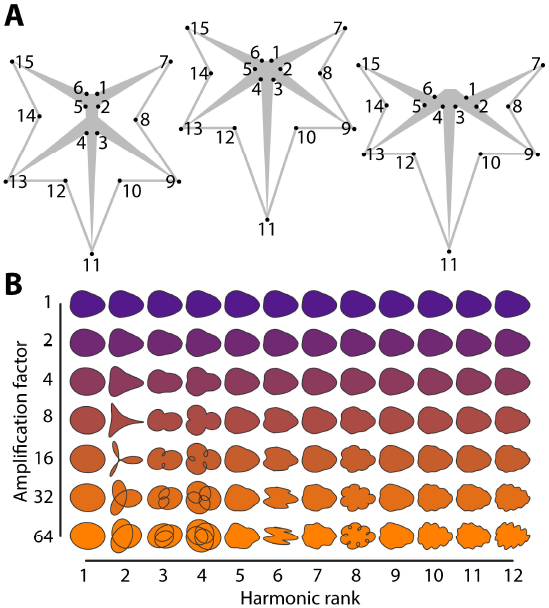
Landmarks and harmonic contributions to shape. **A)** The 15 landmarks used for analysis. Left to right, landmark placement when the distal and proximal veins l) pinnately emerge from the midvein, m) both originate from the petiolar junction, or r) the proximal vein branches from the distal. **B)** Harmonic contributions to shape resulting from Elliptical Fourier Descriptor (EFD) analysis. The harmonic rank is arranged horizontally and the amplification factor (which multiplies the harmonic contributions to shape by the indicated amount) vertically. Note: for convenience to the reader, these panels are recapitulated in the companion manuscript [52].

To determine the extent that landmarks capture qualitative variation in leaf shape among *Passiflora* species, representative leaves were compared to averaged Procrustes-adjusted landmark values (Fig. 2). The landmark analysis captures features such as the relative lengths and angular placement of the proximal and distal veins as well as the depth of the sinuses. Visualizing superimposed landmarks for all leaves measured in addition to the averaged landmark values demonstrates substantial sources of shape variance in some species, especially due to changes in leaf shape across the leaf series, that usually relate to the depth of the sinuses or the number of lobes.

**Figure 2:**
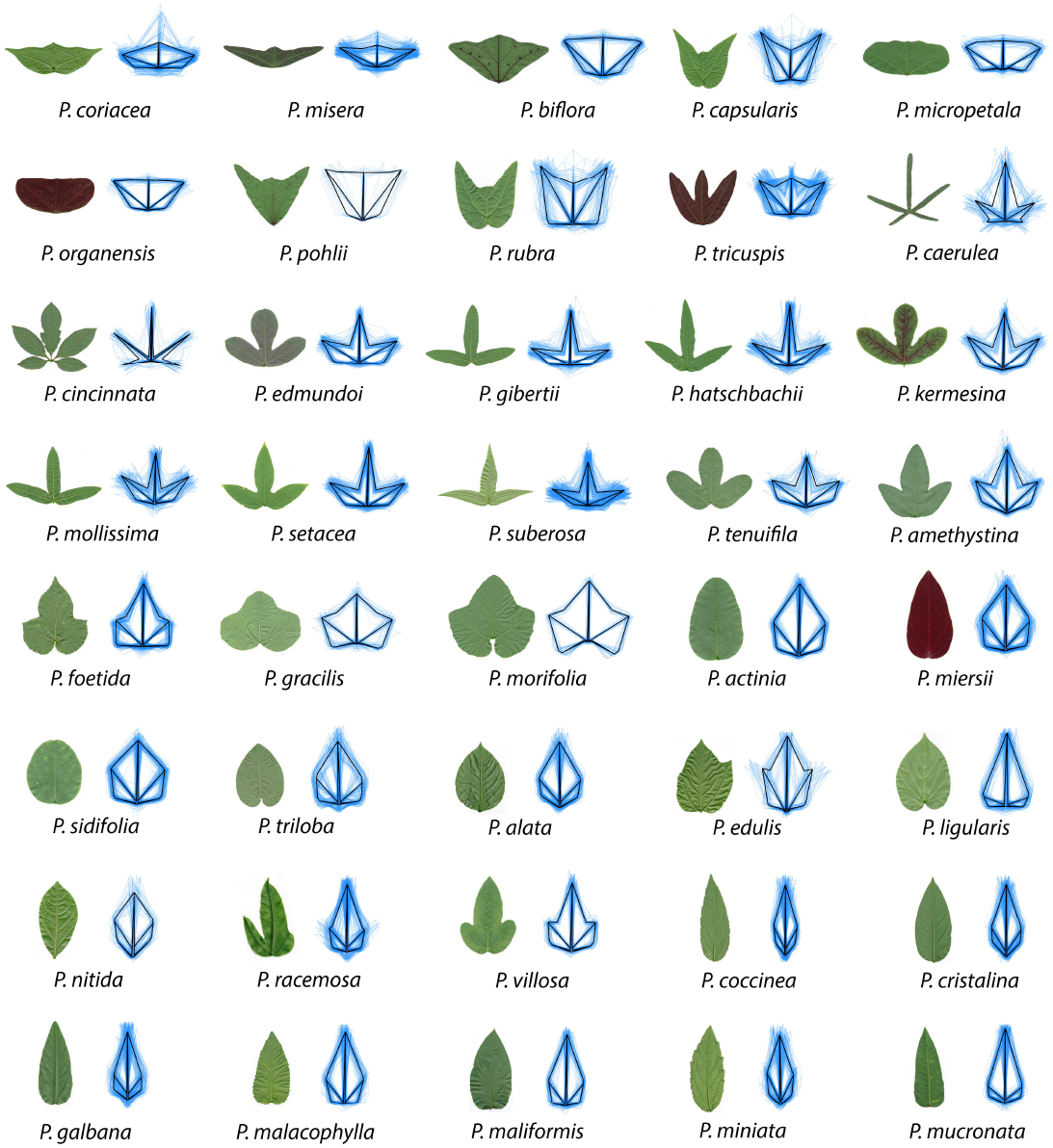
The shapes of *Passiflora* leaves measured using landmarks. For the 40 species analyzed in this study, both a representative leaf and landmark data are shown. For the landmark data, the mean leaf for the species in shown in black, whereas all data for the species is depicted in semi-transparent blue.

Although landmarks accurately depict information related to vascular patterning and the relative placement of the lobes and sinuses of the blade, they fail to capture more subtle shape variation related to the curvature of the lamina. Elliptical Fourier Descriptors (EFDs) result from a harmonic decomposition of a shape outline. The harmonic contributions to leaf shape can be visualized (Fig. 1B), which in *Passiflora* correspond to features reflecting the leaf tip, distal lobes, and proximal lobes (the “trifoliate” features, especially in the lower harmonic ranks) or more local features (the “serrations” represented in the higher harmonic ranks) (Fig. 1B). The averaged outlines of leaves capture the curves and lobing of leaves from each species (Fig. 3). Species that display leaves with variable numbers of lobes (such as *Passiflora caerulea, P. cincinnata,* or *P. suberosa)* have average leaf outlines reflecting this source of shape variance.

**Figure 3:**
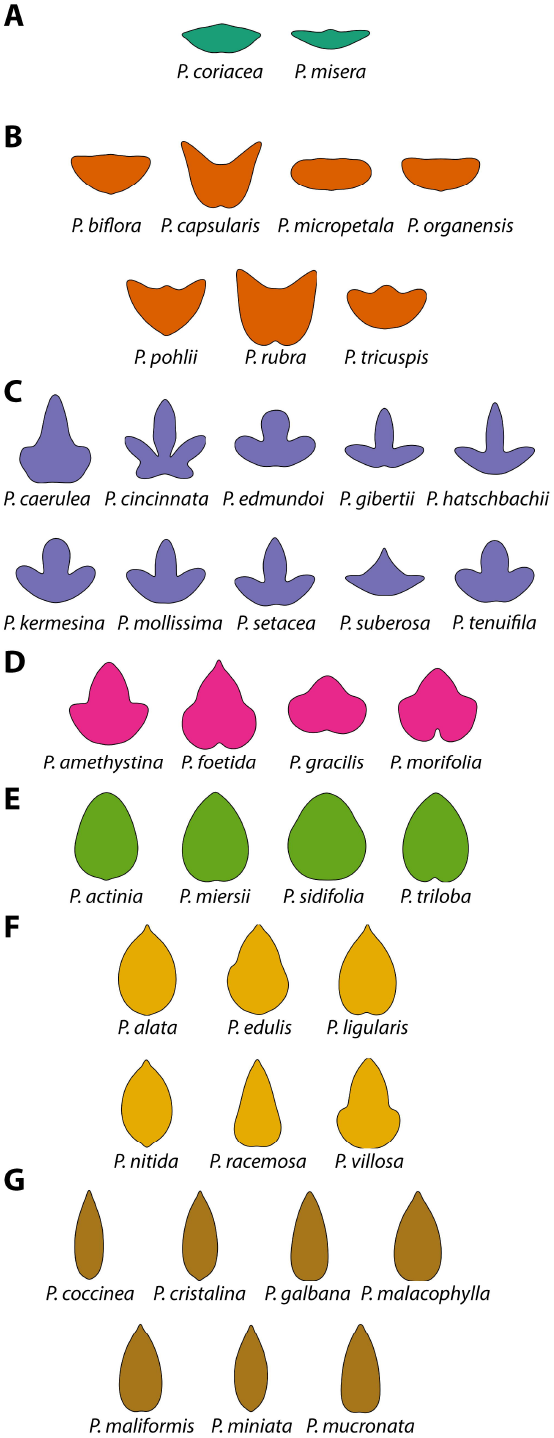
The shapes of *Passiflora* leaves measured using Elliptical Fourier Descriptors (EFDs). Mean leaves calculated for each of 40 species analyzed in this study from the harmonic series resulting from an Elliptical Fourier Descriptor (EFD) analysis of the leaf contours. **A-G)** Classes of species are indicated by their respective panels. Species classes were determined by neighboring position in the Principal Component Analysis (PCA) morphospace, described in Figs. 4**-**5. Color indicates class: class A, teal; class B, orange; class C, lavender; class D, magenta; class E, green; class F, yellow; class G, brown.

### The morphospace reflects species and heteroblastic differences in leaf shape

To analyze major sources of shape variance in Procrustes-adjusted landmark values and the harmonic series from the Elliptical Fourier Descriptor (EFD) analysis, a Principal Component Analysis (PCA) was performed to reduce the dimensionality of each dataset. Onto the resulting morphospaces were projected species identity and the node position in the leaf series (“heteroblasty”). Node position is referred to as “heteroblasty” as a shorthand indicating that numbering of nodes begins at the shoot base, with “1” indicating the first emerged leaf at the shoot base. This numbering scheme more closely aligns with the heteroblastic series of leaves compared to the reverse numbering that begins at the growing shoot tip and is more sensitive to the allometric changes in rapidly expanding leaves.

Eigenleaves (theoretical leaf shapes representing the eigenvectors from a Principal Component Analysis) from each PCA reveal the shape features contributing to shape variance along each Principal Component (PC). The first four landmark PCs (Fig. 4A) explain 83.2% of shape variance for the landmark dataset. PC1 reflects shape variance related to long, lance-like leaves versus wider leaves with short midveins and long, extended distal lobes. Both PC2 and PC3 explain shape variance related to leaves with pronounced distal lobes versus more rounder (PC2) or deltoid (PC3) leaves with less lobing. PC4 also explains shape variance related to lobing. A comparison of the landmark eigenleaves (Fig. 4A) with the EFD eigenleaves (Fig. 4B) shows that the shape variance explained by each respective PC is strikingly similar, especially with respect to lobing and the length-to-width ratio of leaves. This demonstrates a qualitative correspondence between the orthogonal axes of each dataset, including their directionality, which will be subsequently explored in further detail.

**Figure 4:**
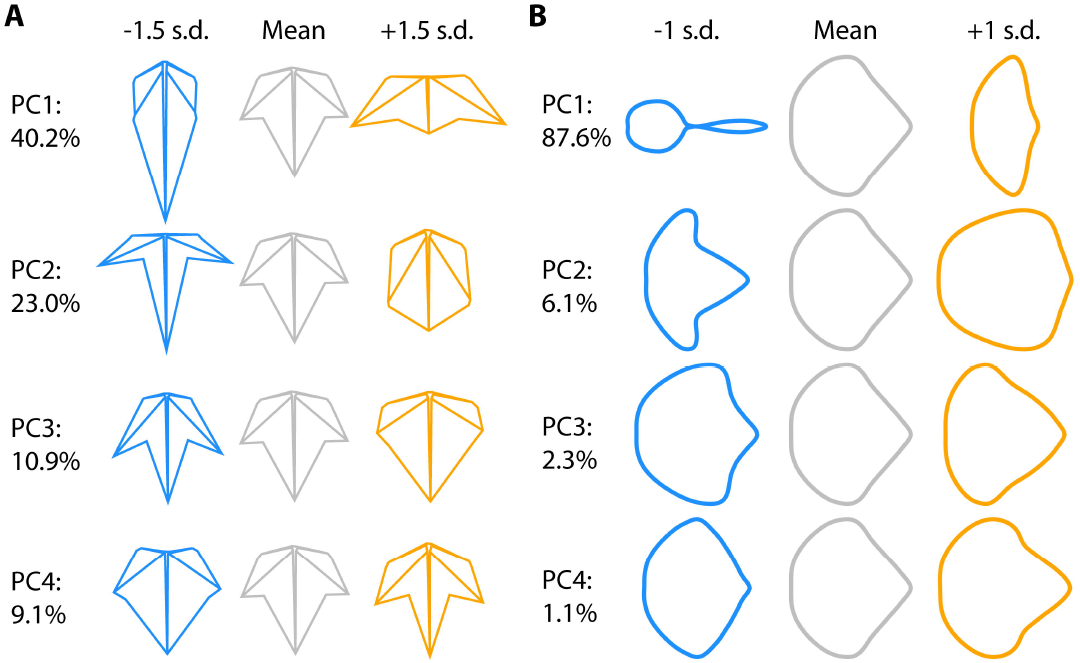
Principal Components (PCs) and eigenleaves. **A)** Principal components (PCs) representing shape variance in landmark data. Eigenleaf representations (theoretical leaf shapes representing the eigenvectors from a Principal Component Analysis) at +/− 1.5 standard deviations (s.d.) are shown for the first four PCs. Percent variance explained by each PC indicated. **B)** PCs representing shape variance in Elliptical Fourier Descriptor (EFD) data. Eigenleaf representations at +/− 1 s.d. are shown for the first four PCs. Percent variance explained by each PC indicated.

Projecting species identity and heteroblastic node onto the landmark and EFD morphospaces reveals that each method separates the shape variance attributable to these variables, but in different ways (Fig. 5). Because visualizing 40 distinct species is a challenge, species were assigned to 7 different classes (consistently colored throughout the manuscript) based on a) occupying similar spaces within morphospace and b) qualitative differences in leaf shape (Fig. 5A). Species classes show pronounced separation from each other by PC1 and PC2 in both the landmark (Fig. 5B) and EFD (Fig. 5C) morphospaces. Less separation is observed by species class for PC3 and PC4. When heteroblastic node is projected onto the morphospaces, there is a trend for the leaves originating from high heteroblastic nodes (young leaves towards the growing tip) to occupy the lower PC2 values within each species class. This is especially true for the landmark morphospace (Fig. 5B). There is also a trend for leaves originating from high heteroblastic nodes to have low PC3 values, regardless of species class. Both low PC2 and PC3 values correspond to more pronounced distal lobing (Fig. 4), a shape feature commonly found in young leaves near the growing tip of the plant, compared to older leaves near the base of the vine that tend to have less lobing. That shape variance attributable to species class and heteroblastic node traverse the morphospace in different ways suggests to some extent the shape variance for each of these factors is separable, as is discussed in the next section.

**Figure 5:**
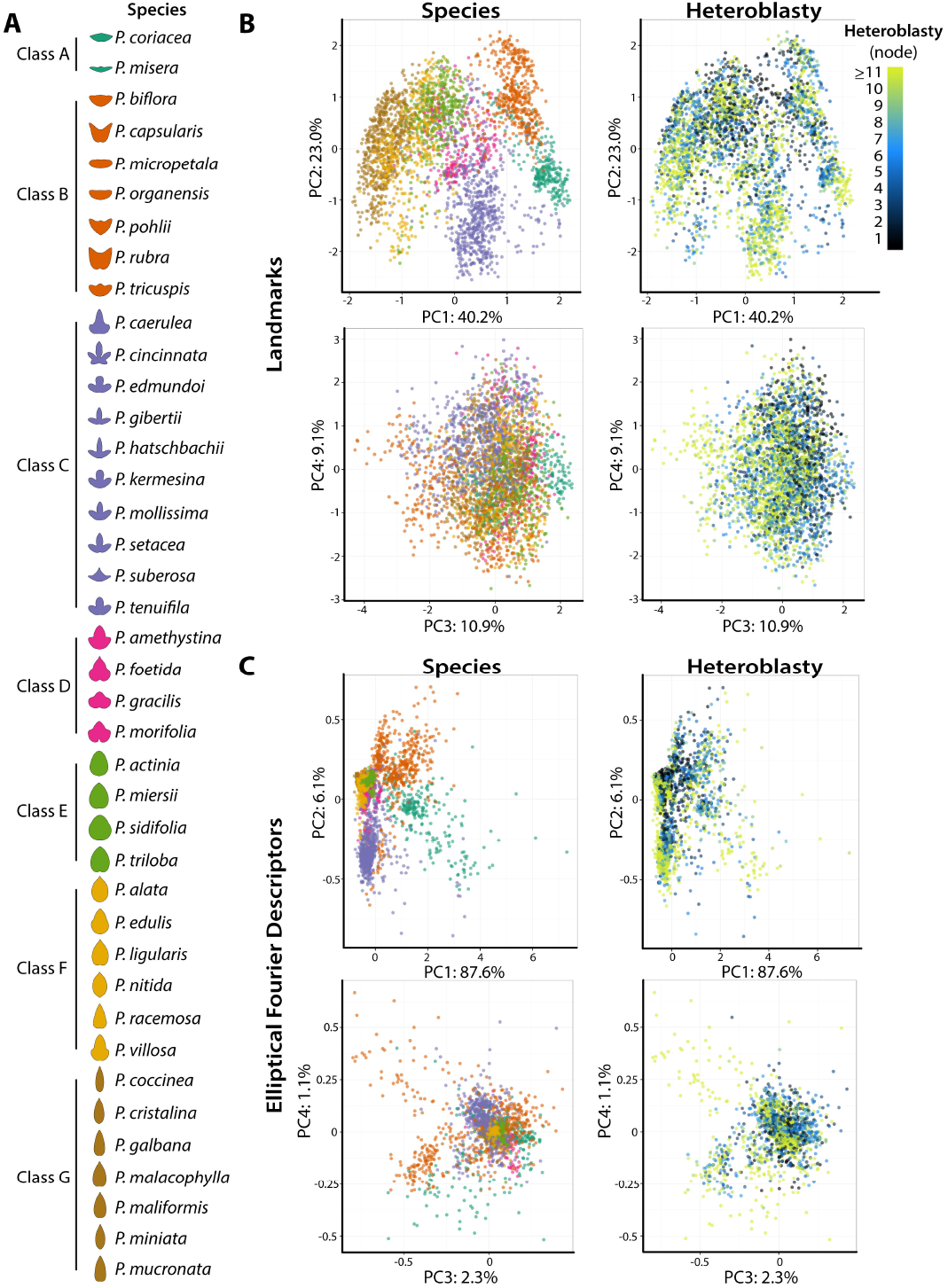
Morphospace by species and heteroblastic node. **A)** Key, showing species classes and averaged leaf contours for each species. Color indicates class, which is used in other panels. **B)** Principal Component Analysis (PCA) of landmark data. Graphs for PC2 vs. PC1 and PC4 vs. PC3 are colored by species class and by heteroblastic node. Percent variance explained by each PC indicated. **C)** PCA of Elliptical Fourier Descriptor (EFD) data. Graphs for PC2 vs. PC1 and PC4 vs. PC3 are colored by species class and by heteroblastic node. Percent variance explained by each PC indicated. Heteroblastic node position is numbered “1” starting from the shoot base. Class color scheme: class A, teal; class B, orange; class C, lavender; class D, magenta; class E, green; class F, yellow; class G, brown. Heteroblastic node color scheme: shoot base, black; middle shoot, blue; shoot tip, yellow.

### Discriminating species vs. node identity

That species class and node identity traverse the morphospace differently (Fig, 5A-B) is consistent with previous work demonstrating that shape features can be used to discriminate species independently from node position in grapevine [10, 24]. A Linear Discriminant Analysis (LDA) is used here to determine the extent these two variables can be predicted independently of the other in *Passiflora* using landmark data, Elliptical Fourier Descriptors (EFDs), and both landmark and EFDs together. We stress that the LDA approach taken in this work is fundamentally different from modeling species, node, and interaction effects using linear modeling. Such an approach (which we undertook but the data is not shown here, because it is outside the scope of this manuscript) reveals that for each morphometric trait considered independently, the species and interaction effects are the strongest and the node effect is weak. Rather, an LDA allows explicit questions to be asked regarding all the measured traits together. Can all the traits be used together to discriminate species regardless of node? Using all traits can node be distinguished separately from species? Such a framework is consistent with developmental genetic theory that differences in leaf shape between species versus more conserved heteroblastic changes in leaf shape within individual plants are regulated by distinct genetic pathways [16] that lead to separable morphological effects within single leaves (so called “cryptotypes” [46]). We also note that the LDAs performed use the “leave one out” approach of cross-validation, in which a separate LDA for each leaf, minus the leaf in question, is used to predict the identity of that leaf. Such an approach is designed to compensate for differences in species replication and nodes sampled per a vine in our dataset (see raw data [45] and **Fig. S1**).

An LDA is first performed on species identity, regardless of node position. The resulting discriminants are then used to predict the identity of the species. Regardless of whether landmarks (Fig. 6A), EFDs (Fig. 6C), or both landmarks and EFDs are used (Fig. 6E) a high proportion of leaves can be correctly reassigned to the correct species. When there is confusion between species, it tends to be within the same species class. This result demonstrates that regardless of the position of a leaf within the heteroblastic series, its species identity can be predicted. For most species classes (all except C and D) the maximum correct prediction is most often achieved with both landmark and EFD data together compared to each data type alone (Table 1). For species classes C and D, however, landmark data alone tends to outperform EFD and both data types together. This indicates that for some species, especially those that are highly lobed as in species classes C and D, landmark data is a better indicator of species identity (perhaps because it is more explicitly related to lobing).

**Figure 6:**
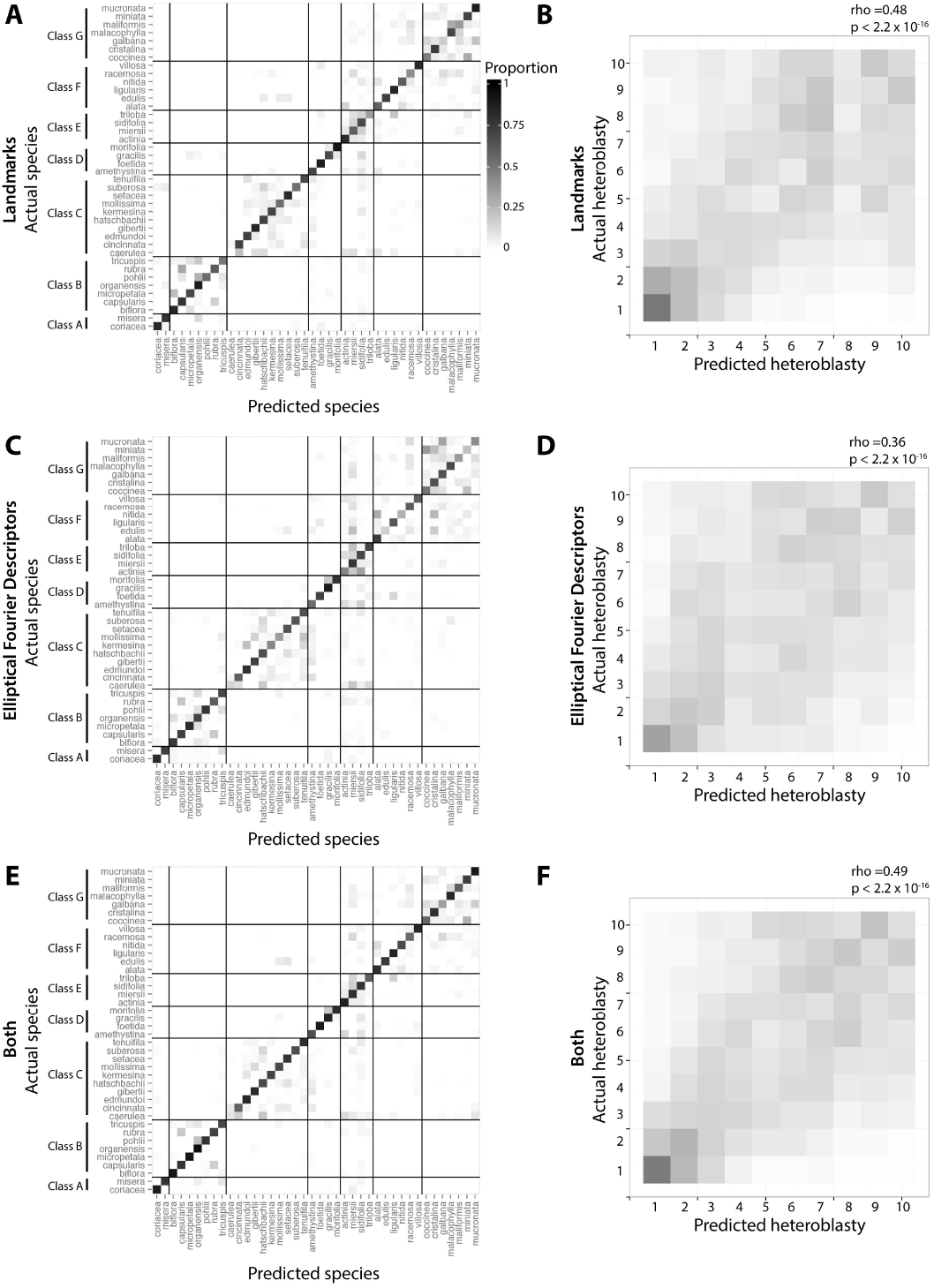
Linear Discriminant Analysis (LDA). Linear Discriminant Analysis (LDA) using **A-B)** landmark data, **C-D)** Elliptical Fourier Descriptor (EFD) data, and **E-F)** both datasets. For each set of LDAs, analysis was performed to discriminate species (ignoring heteroblastic node information) or to discriminate heteroblastic node (ignoring species information). Subsequent prediction of species or heteroblastic node identity is then visualized using confusion matrices, where actual identity is oriented vertically, predicted identity horizontally, and the proportion assigned indicated as fill. Species LDAs are broken up by species class. For heteroblastic node LDAs, Spearman’s rho and associated p values calculated from correlating actual and predicted node identities are provided. Predictions carried out using LDA use the “leave one out” approach cross-validation approach. Heteroblastic node position is numbered “1” starting from the shoot base. Color scheme: low assigned proportion, white; high assigned proportion, black.

**Table 1:**
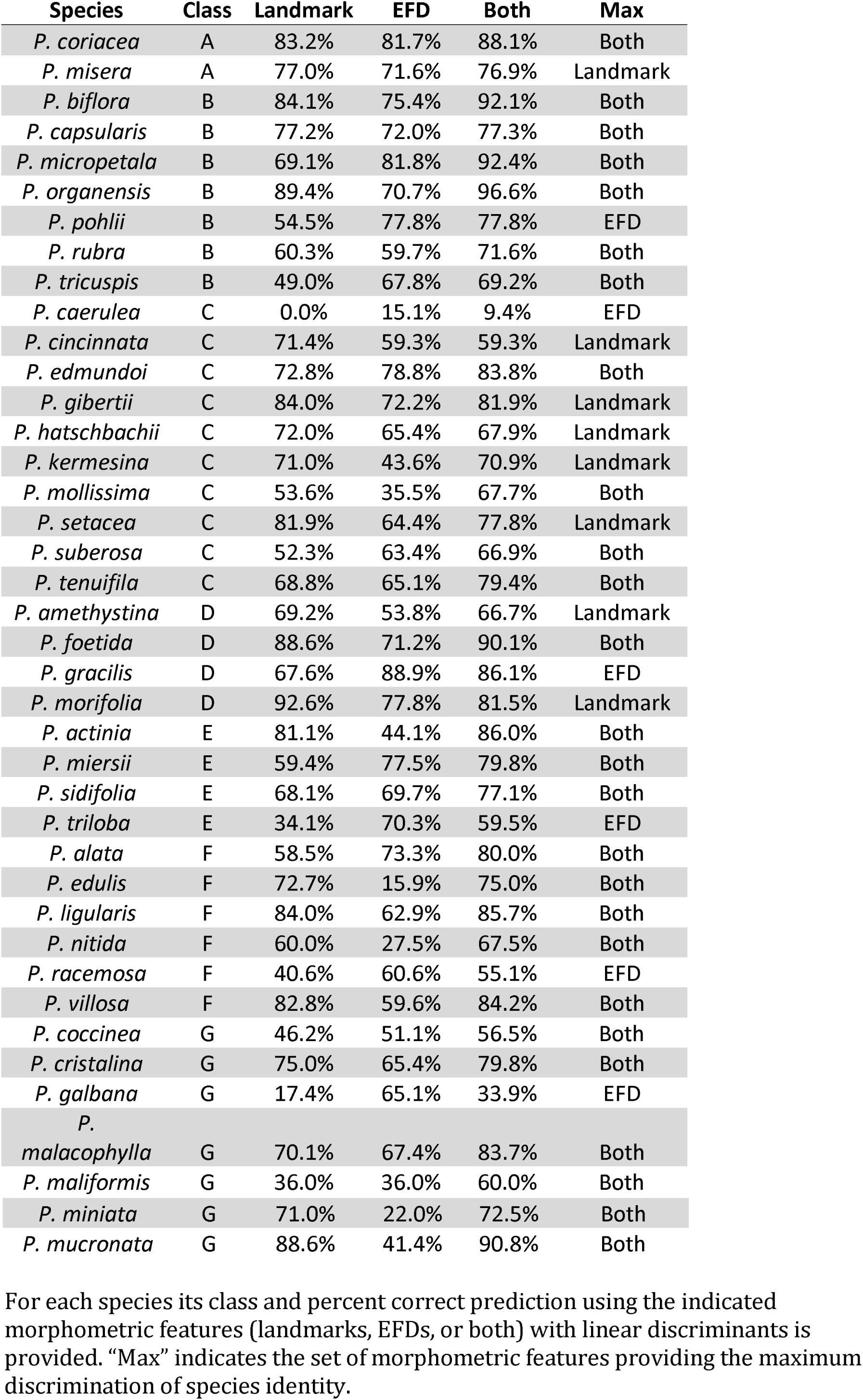
Predictive power of different morphometric methods to discriminate *Passiflora* species.

Conversely, heteroblastic node position can be predicted independently of species identity, but to a much lesser degree and not equally across the leaf series. The leaves occupying lower node positions (older leaves at the base of the vine) tend to be successfully predicted at a higher rate than the younger leaves of the tip, regardless of whether landmarks (Fig. 6B), EFDs (Fig. 6D), or both landmarks and EFDS are used (Fig. 6F). EFDs, however, overall under-perform landmarks or landmarks and EFDs used together (Table 2). This indicates that landmarks are a superior discriminant of node position compared to EFDs. Previous work in grapevine indicates that vein thickness is altered by shoot position [10, 24]. That landmarks measure vein thickness, but not EFDs, may explain the differing abilities of these two shape features to correctly discriminate leaves by heteroblastic node position. That the juvenile leaves at the lower heteroblastic node positions are correctly predicted at higher rates suggests that these leaves are more similar across species (or correspondingly, that leaves at high heteroblastic node positions are more divergent between species).

**Table 2:**
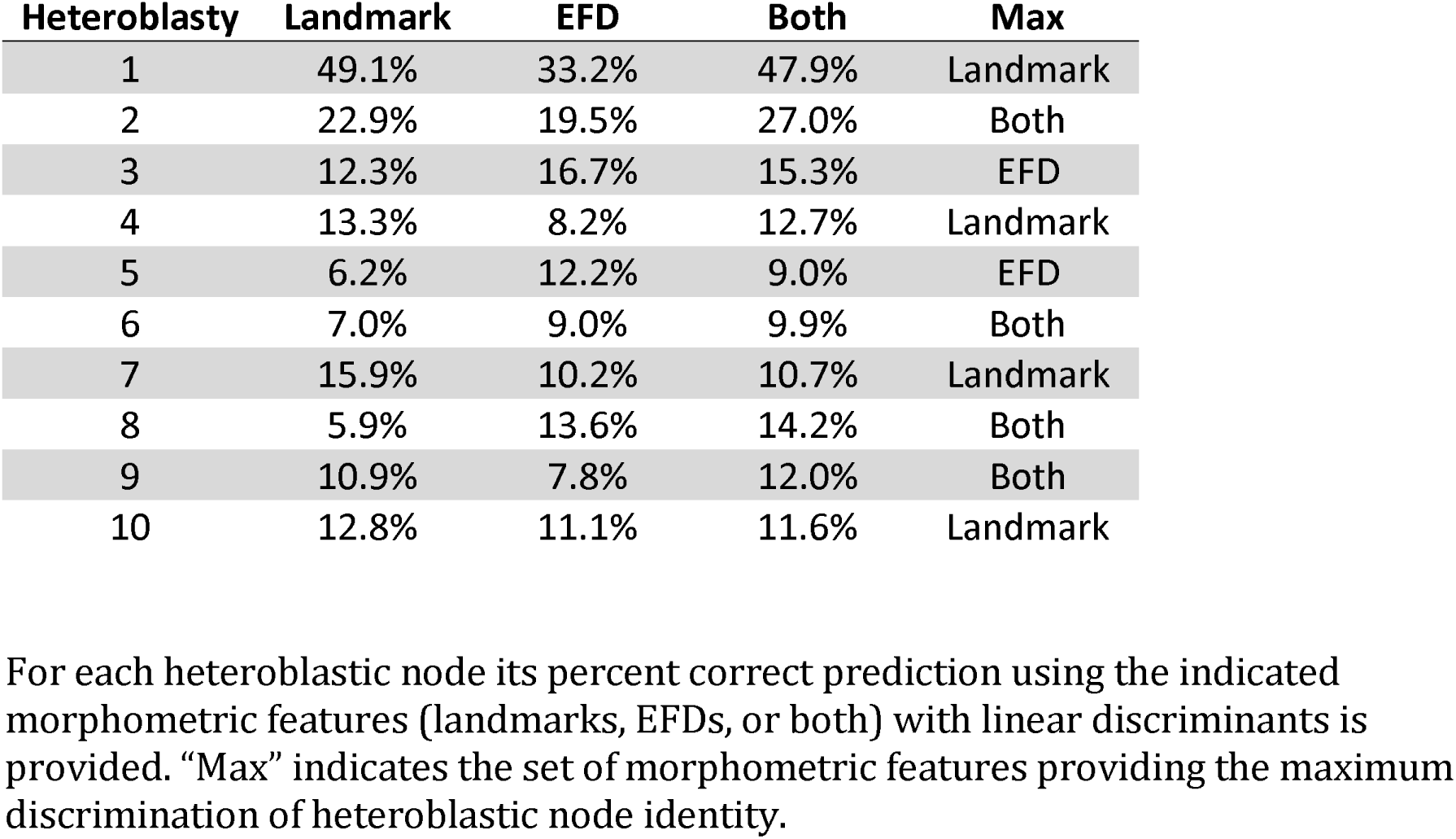
Predictive power of different morphometric methods to discriminate heteroblastic node.

| Heteroblasty | Landmark | EFD | Both | Max |
| --- | --- | --- | --- | --- |
| 1 | 49.1% | 33.2% | 47.9% | Landmark |
| 2 | 22.9% | 19.5% | 27.0% | Both |
| 3 | 12.3% | 16.7% | 15.3% | EFD |
| 4 | 13.3% | 8.2% | 12.7% | Landmark |
| 5 | 6.2% | 12.2% | 9.0% | EFD |
| 6 | 7.0% | 9.0% | 9.9% | Both |
| 7 | 15.9% | 10.2% | 10.7% | Landmark |
| 8 | 5.9% | 13.6% | 14.2% | Both |
| 9 | 10.9% | 7.8% | 12.0% | Both |
| 10 | 12.8% | 11.1% | 11.6% | Landmark |

For each heteroblastic node its percent correct prediction using the indicated morphometric features (landmarks, EFDs, or both) with linear discriminants is provided. “Max” indicates the set of morphometric features providing the maximum discrimination of heteroblastic node identity.

### Correlational matrix between landmarks and Elliptical Fourier Descirptors (EFDs)

Until now, landmarks and Elliptical Fourier Descriptors (EFDs) have either been considered separately or in conjunction together but not compared against each other. The landmarks used in this study tend to represent vascular features of the leaf, the lobes, and the sinuses. The EFDs represent the blade and the continuous contour and curves of lamina. Further, landmark data is represented as (x, y) coordinates, whereas EFD data is a Fourier-based harmonic series. A correlational matrix is used to find strong associations between the components of each dataset and to interpret the features each dataset uniquely quantifies against the other.

The input for the correlation matrix, using Spearman’s rho, is each of the fifteen x and y coordinates of the landmark dataset and each of the four harmonic coefficients (A, B, C, D) of the first 20 harmonic ranks from the EFD data, correlated across the >3,300 leaves for all species and heteroblastic node positions used in this study. This correlation matrix was used as a distance matrix to hierarchically cluster these traits and the rho and p values subsequently visualized (Fig. 7).

**Figure 7:**
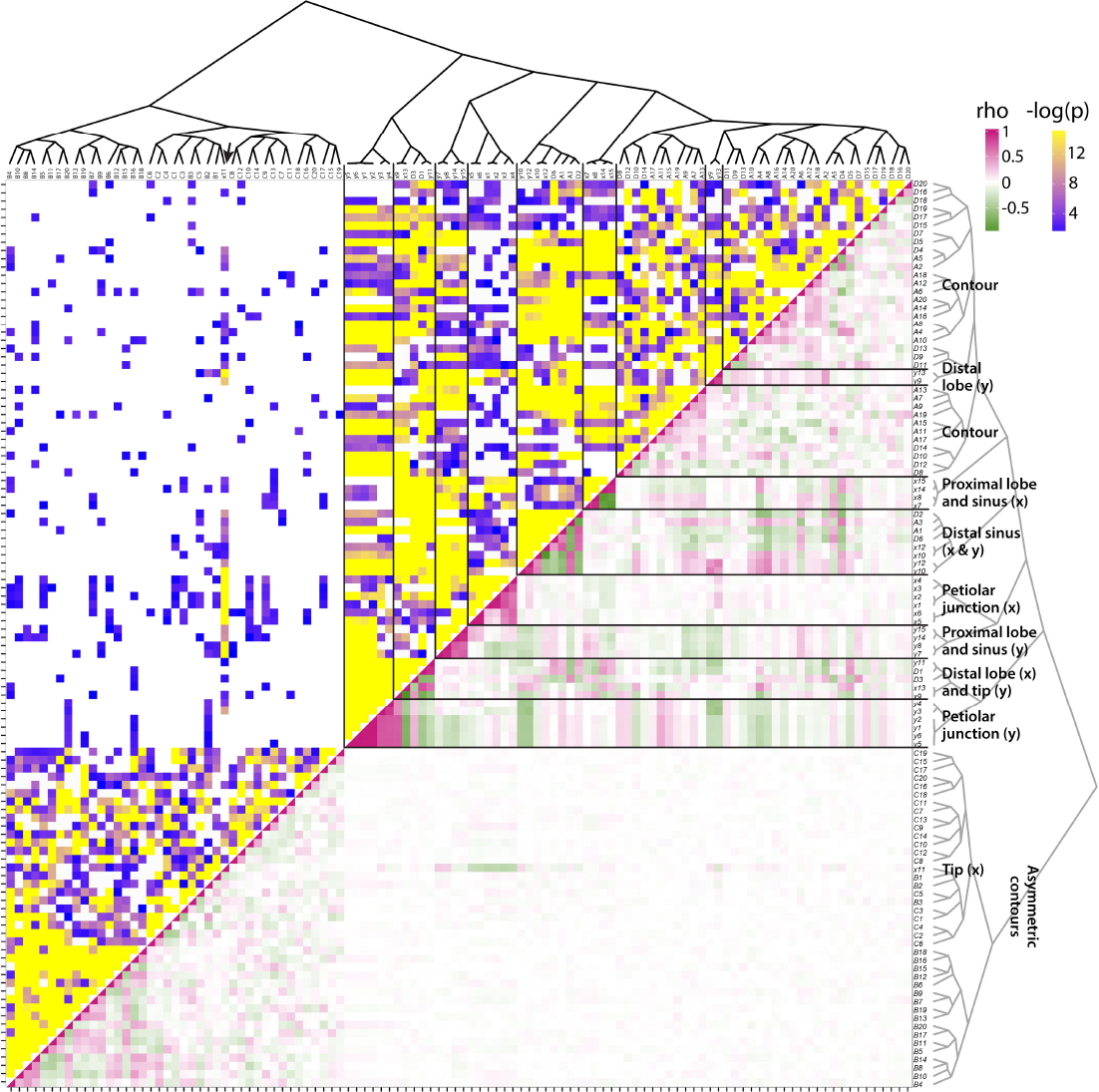
Correlational matrix of landmark and Elliptical Fourier Descriptor (EFD) traits. Spearman’s correlation matrix for morphometric features analyzed in this study. Upper half indicates –log_10_ p value and lower half Spearman’s rho between indicated traits. Morphometric traits, both landmark and the harmonic series, are indicated along the sides, arranged using hierarchical clustering, the topology of which is depicted as a dendrogram. Key groupings of landmarks indicating correlational associations with each other or EFD harmonics are indicated. Spearman’s rho: low values, green; middle values, white; high values, magenta. −log_10_ p values: low values, purple; high values, yellow; p < 0.05, no color.

A large set of uncorrelated traits, consisting of the B and C harmonic coefficients and the x11 landmark, end up clustered together (Fig. 7). The B and C harmonic coefficients represent asymmetric sources of shape variance [31] and the x11 landmark represents the left-right variance of the leaf tip (Fig. 1A), which will mostly be affected by leaf asymmetry. That these shape features are weakly correlated with each other and other traits only implies that they are regulated by an unaccounted source of variance for this particular analysis. In the future, a more in-depth analysis will likely reveal phyllotaxy as modulating leaf asymmetry [39, 40], specifically alternating asymmetry at consecutive nodes, as recently shown in other vines, such as ivy and grapevine [25].

The remaining landmarks and the A and D coefficients of the harmonic series (representing symmetrical shape variation) show various correlational associations with each other (Figs. 7**-**8). Harmonic contributions to leaf shape (Fig. 1B) are more abstract and difficult to interpret than the contributions of landmarks to leaf shape, as the landmarks represent homologous points found in every leaf (Fig. 1A). Strong correlations between harmonic coefficients with landmarks can help interpret the context of the harmonic coefficient to leaf shape. Most the harmonic coefficients cluster exclusively together except for landmarks y9 and y13, which represent the proximal-distal displacement of the distal lobes along the leaf length (Fig. 8). This suggests that large amounts of the shape variance associated with the contour of the blade are influenced by the relative placement of the distal lobes along the leaf length. The remaining harmonic coefficients that cluster outside most the other coefficients also associate with features of the distal part of the leaf. A1, A3, D2, and D6 associate with the x and y coordinates of the distal sinus (x10, x12, y10, and y12) and D1 and D3 associate with the left-right displacement of the distal lobe (x9 and x13) and the vertical displacement of the leaf tip (y11) (Fig. 8). Although difficult to interpret, the correlations of harmonic coefficients suggest that the overall leaf contour is influenced by the placement of the distal lobe and sinus.

**Figure 8:**
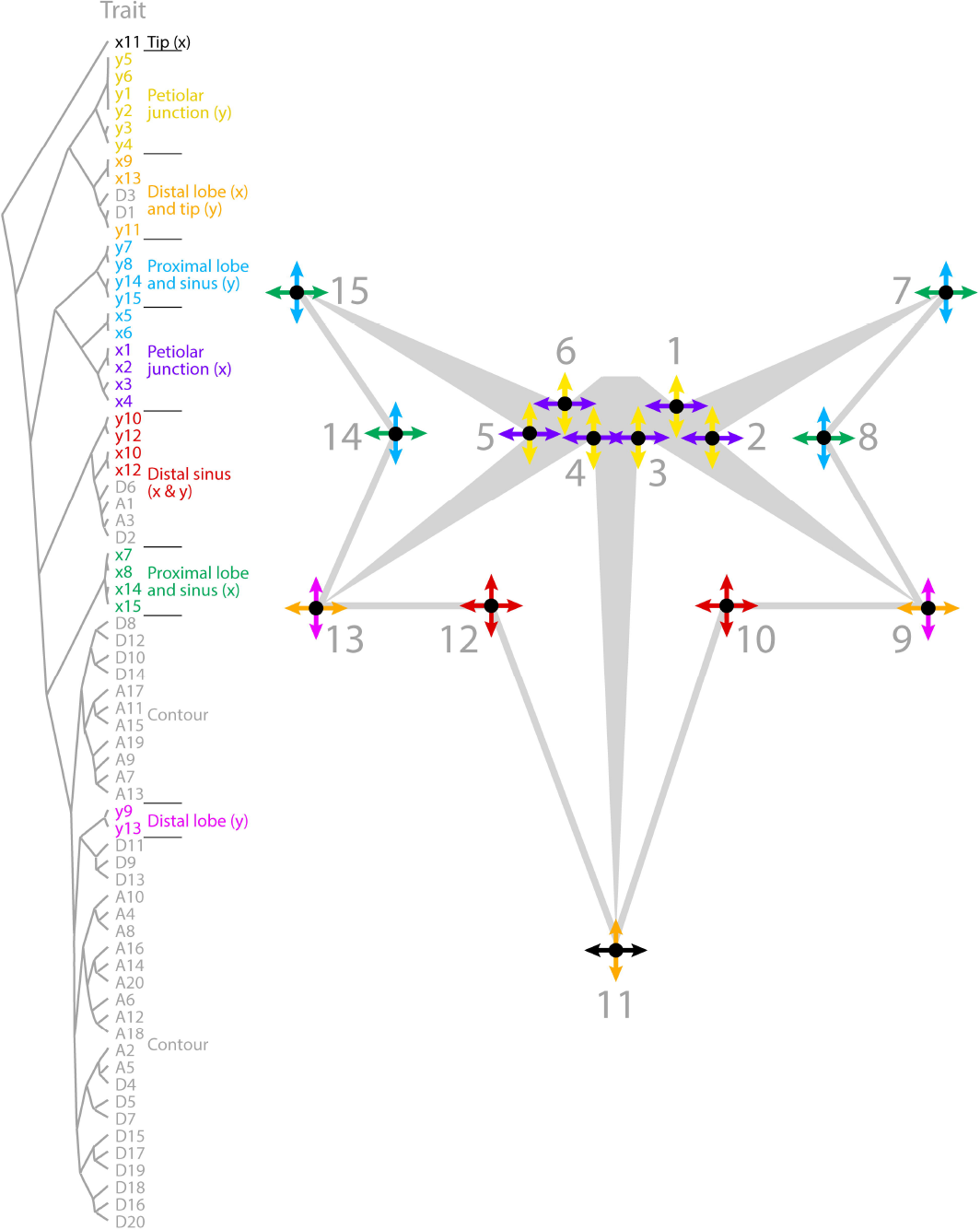
Correlational relationships between vascular landmarks and leaf contours. Correlational relationships between x and y components of landmarks and Elliptical Fourier Descriptor (EFD) harmonics are indicated by dendrogram (left) and landmarks qualitatively on a representation of a leaf (right). x and y landmark components are independently depicted by arrows and colored as indicated to show major correlational sources of shape variance within *Passiflora* leaves.

The remaining correlations between landmarks reveal interesting constraints governing the shape of *Passiflora* leaves (Fig. 8). As mentioned previously, the left-right displacement of the distal lobes (x9 and x13) strongly correlates with the vertical proximal-distal displacement of the leaf tip (y11). The x and y coordinates of the distal lobes (landmarks 10 and 12) are the only features for which the x and y displacement are correlated, suggesting that the distal sinus varies in a diagonal direction. The proximal sinus and lobe (landmarks 7, 8, 14, and 15) and the landmarks at the base of the veins of the petiolar junction (landmarks 1, 2, 3, 4, 5, and 6) form additional groups of associated landmarks, although interestingly the x and y displacement of each of these two groups is distinct in each case (Fig. 8)

## Discussion

Leaf morphology refers to the totality of leaf architecture, at the cellular, tissue, and organ levels, and distinct attributes of the leaf, both the vasculature and lamina. The topology of the vasculature and contour of the leaf blade are distinct geometric phenomena that require different morphometric approaches to quantify. Landmarks and Elliptical Fourier Descriptors (EFDs) are ideal methods to analyze the distinct features of leaves contributing to their shape (Fig. 1), but rarely are they measured and compared on the same leaves. Our analysis of disparate leaf shapes among *Passiflora* species with landmarks (Fig. 2) and EFDs (Fig. 3) reveals that both methods capture similar orthogonal axes of shape variation (Fig. 4), and separate both species and heteroblastic node identity, but in distinct ways (Fig. 5). Landmarks are superior to EFDs in predicting node position compared to species identity, most likely because they describe vascular patterning, which is relatively sensitive to heteroblasty compared to species differences in leaf shape (Fig. 6; Tables 1–2). Although most elements of the EFD harmonic series cluster together in a pairwise correlational analysis, a few are closely associated with landmarks (Fig. 7). Landmarks exhibit a correlational structure revealing developmental constraints in how leaves vary across *Passiflora* species and the heteroblastic series (Fig. 8). Together, our data quantify the relationship between blade and vasculature, revealing that one does not drive the patterning of the other, and although each distinctly varies, many shape features of the leaf change in concert across evolution and development.

## Methods

### Plant materials and growth conditions

*Passiflora* germplasm was kindly provided by R. Silva (Viveiros Flora Brasil, Araguari, MG, Brazil), Dr. F.G. Faleiro (EMBRAPA Cerrados, Planaltina, DF, Brazil), Prof. M.M. Souza (Universidade Estadual de Santa Cruz - UESC, Ilhéus, BA, Brazil), M. Peixoto (Mogi das Cruzes, SP, Brazil), Prof. M.L. Silva (Universidade do Estado de Mato Grosso, Tangará da Serra, MT, Brazil), and Prof. C.H. Bruckner (Universidade Federal de Viçosa, Viçosa, MG, Brazil).

The plants were germinated from seed, planted between late October 2015 and early March 2016, in Viçosa, at the Federal University of Viçosa, MG, Brazil. The populations were raised and maintained under polycarbonate-covered greenhouse conditions, equipped with automatic environmental control using exhaust fans and evaporative cooling panels (with expanded clay wettable pads). Seeds for each *Passiflora* species were sown in 128 cell propagation plastic trays (GPlan Comércio de Produtos Agrícola s EIRELI – ME, São Paulo, SP, Brazil) filled with horticultural organic Tropstrato HT Hortaliças substrate (Vida Verde Indústria e Comércio de Insumos Orgânicos Ltda, Mogi Mirim, SP, Brazil). After germination (30-40 days), plantlets were individually transplanted to 5 L capacity plastic pots (EME-A-EME Ind. Com. Ltda., Petrópolis, RJ, Brazil) filled with horticultural substrate. Each pot received 5 g of Osmocote^®^ Plus NPK 15-09-12 3-4 month controlled release fertilizer (Scotts, USA). Plants were irrigated on a daily-basis with tap water, and no phytosanitary control was applied. The germination and growth rates of plants varied widely. The number of replicates for each species and the number of nodes per vine are indicated in the raw data [45] and depicted visually (**Fig. S1**).

For scanning, a multifunction printer (Canon PIXMA MX340 Wireless Office All-in-One Printer, model 4204B019, USA) was used. A 20 cm metallic ruler was positioned at the bottom of each scanned sheet as a size marker. Leaves were carefully detached, from the base to the tip of the shoot, and affixed to an A4 paper sheet, adaxial face down, using 12 mm-double sided tape (Scotch Model 9400, 3M do Brasil, SP, Brazil). The numbers written near each leaf indicate position in the shoot, in a tip-to-base direction, starting with the youngest leaf at the tip of the shoot. It should be noted that the numbering in the scans is opposite from the numbering used in the analysis and figures for this manuscript, in which leaves are numbered with “1” starting at the shoot base. This numbering system more closely aligns with the heteroblastic series than the reverse numbering scheme originally used in the scans.

### Morphometric and statistical analyses

All morphometric data and code used for statistical analysis is available on GitHub [45]. All original data is available at GigaDB [44].

Landmarks, as described in the text, were placed on leaves in ImageJ [47]. Procrustes superimposition was performed using the shapes package [48] in R [49] with the procGPA function using reflect=TRUE. Resulting Procrustes-adjusted coordinates and principal component scores (PCs) were written out for subsequent analyses and eigenleaf representations visualized using the shapepca function.

To isolate outlines for Elliptical Fourier Descriptor (EFD) analysis, the “Make Binary” function in ImageJ [47] was found to be sufficient to segment leaves. The wand tool was used to select individual binary leaf outlines, which were pasted into a new canvas, which was subsequently saved as an individual image, which was named by vine and node position from which the leaf was derived. The binary images were batch converted into RGB .bmp files and read into SHAPE, which was used to perform chain-code analysis [31, 32]. The resulting chain-code .chc file was then used to calculate normalized EFDs. The resulting normalized EFD .nef file was then read into Momocs (version 0.2-6) [33] in R. The harmonic contributions to shape were visualized using the hcontrib function. Averaged leaf outlines were calculated using the meanShapes function and Principal Component Analysis (PCA) performed using the pca function and eigenleaves visualized using the PC.contrib function.

Unless otherwise noted, all visualization was performed using ggplot2 in R [50]. Linear Discriminant Analysis (LDA) was performed using the lda function and subsequent prediction of species identity or heteroblastic node position performed using the predict function with MASS [51]. When LDAs were used for prediction, the parameter CV was set to “TRUE”, for the “leave one out” cross-validation approach, to help make analyses more robust to differences in replication and node numbers between species and vines. Hierarchical clustering was performed using the hclust function.

## Availability and requirements

Project name: PassifloraLeaves

Project home page: https://github.com/DanChitwood/PassifloraLeaves

Operating system(s): Platform independent

Programming language: R

Other requirements: Not applicable

License: MIT license

Any restriction to use by non-academics: none

## Availability of supporting data and materials

The data sets supporting the results of this article are available in the GigaDB repository [44].

## Declarations

### Funding

Brazilian sponsoring agencies, namely FAPEMIG (Grant no. CBB - APQ-01131-15), CNPq (Grant no. 459.529/2014-5) and CAPES, are acknowledged for financial support.

## Authors’ contributions

The overall project was conceived by DHC and WCO. WCO grew and scanned all plant material and DHC carried out analysis. DHC and WCO wrote the paper.

## Competing interests

The authors declare that they have no competing interests.

## Supplemental Information

**Figure S1: Species replication and number of nodes sampled. A)** Dotplot showing the number of vines sampled for each species. **B)** Boxplot showing the number of nodes sampled for vines from each species. The largest red line is the median nodes sampled (14 nodes), the medium sized redlines the 25^th^ and 75^th^ quantiles (12 and 16 nodes, respectively), and the thin red lines the minimum and maximum (7 and 28 nodes, respectively).

